# Ultrafast glutamate sensors resolve high-frequency release at Schaffer collateral synapses

**DOI:** 10.1101/233494

**Authors:** Nordine Helassa, Céline D. Dürst, Catherine Coates, Silke Kerruth, Urwa Arif, Christian Schulze, J. Simon Wiegert, Michael Geeves, Thomas G. Oertner, Katalin Török

## Abstract

Glutamatergic synapses display a rich repertoire of plasticity mechanisms on many different time scales, involving dynamic changes in the efficacy of transmitter release as well as changes in the number and function of postsynaptic glutamate receptors. The genetically encoded glutamate sensor iGluSnFR enables visualization of glutamate release from presynaptic terminals at frequencies up to ∼10 Hz. However, to resolve glutamate dynamics during high frequency bursts, faster indicators are required. Here we report the development of fast (iGlu_*f*_) and ultrafast (iGlu_*u*_) variants with comparable brightness, but increased *K*_d_ for glutamate (137 μM and 600 μM, respectively). Compared to iGluSnFR, iGlu_*u*_ has a 6-fold faster dissociation rate *in vitro* and 5-fold faster kinetics in synapses. Fitting a three-state model to kinetic data, we identify the large conformational change after glutamate binding as the rate-limiting step. In rat hippocampal slice culture stimulated at 100 Hz, we find that iGlu_*u*_ is sufficiently fast to resolve individual glutamate release events, revealing that glutamate is rapidly cleared from the synaptic cleft. Depression of iGlu_*u*_ responses during 100 Hz trains correlates with depression of postsynaptic EPSPs, indicating that depression during high frequency stimulation is purely presynaptic in origin. At individual boutons, the recovery from depression could be predicted from the amount of glutamate released on the second pulse (paired pulse facilitation/depression), demonstrating differential frequency-dependent filtering of spike trains at Schaffer collateral boutons.

**Significance Statement:** Excitatory synapses convert presynaptic action potentials into chemical signals that are sensed by postsynaptic glutamate receptors. To eavesdrop on synaptic transmission, genetically encoded fluorescent sensors for glutamate have been developed. However, even the best available sensors lag behind the very fast glutamate dynamics in the synaptic cleft. Here we report the development of an ultrafast genetically encoded glutamate sensor, iGlu_*u*_, which allowed us to image glutamate clearance and synaptic depression during 100 Hz spike trains. We found that only boutons showing paired-pulse facilitation were able to rapidly recover from depression. Thus, presynaptic boutons act as frequency-specific filters to transmit select features of the spike train to specific postsynaptic cells.

## INTRODUCTION

The efficacy of synaptic transmission is not constant, but changes dynamically during high-frequency activity. In terms of information processing, different forms of short-term plasticity act as specific frequency filters: facilitating synapses are most effective during high frequency bursts, while depressing synapses preferentially transmit isolated spikes preceded by silent periods (1). Mechanistically, a number of pre- and postsynaptic parameters change dynamically during high frequency activity, e.g. the number of readily releasable vesicles, presynaptic Ca_2+_ dynamics, and the properties of postsynaptic receptors, which may be altered by Ca_2+_-activated enzymes (2, 3).

Electrophysiological analysis of short-term plasticity, by monitoring postsynaptic responses, is complicated by the fact that neurons are often connected by more than one synapse. In addition, it is not straightforward to distinguish between pre-and postsynaptic plasticity mechanisms. Directly measuring glutamate concentrations inside the synaptic cleft during high-frequency activity would allow isolating the dynamics of the vesicle release machinery from potential changes in glutamate receptor properties (e.g. desensitization, phosphorylation or lateral diffusion). Early fluorescent glutamate sensors, constructed by chemical labelling of the fused glutamate binding lobes of ionotropic glutamate receptor GluA2 (termed S1S2) (4-6) and later of the bacterial periplasmic glutamate/aspartate binding protein (GluBP) (7, 8), were not suitable for quantitative single-synapse experiments due to their low dynamic range. Genetically encoded FRET-based fluorescent glutamate sensors e.g. FLIPE, GluSnFR and SuperGluSnFR (**Fig. 1a**) have relatively low FRET efficiency, since glutamate binding causes only a small conformational change in GluBP (9-11). A breakthrough in visualizing glutamate release in intact tissue was achieved with iGluSnFR, a single-fluorophore glutamate sensor (12). Following the concept developed for the GCaMP family of genetically encoded Ca_2+_ sensors (13), iGluSnFR was constructed from circularly permuted (cp) EGFP (14) inserted into the GluBP sequence, creating a large fragment iGlu_l_ (residues 1-253) at the N-terminus and a small fragment iGlu_s_ (residues 254-279) at the C-terminus (**Fig. 1a**). Upon glutamate binding GluBP is reconstituted from its two fragments, pulling the cpEGFP β-barrel together, resulting in a ∼5-fold fluorescence increase. iGluSnFR is targeted for extracellular expression, like previous genetically encoded glutamate sensors, by fusion with a PDGFR peptide segment (10, 12).

**FIGURE 1.**
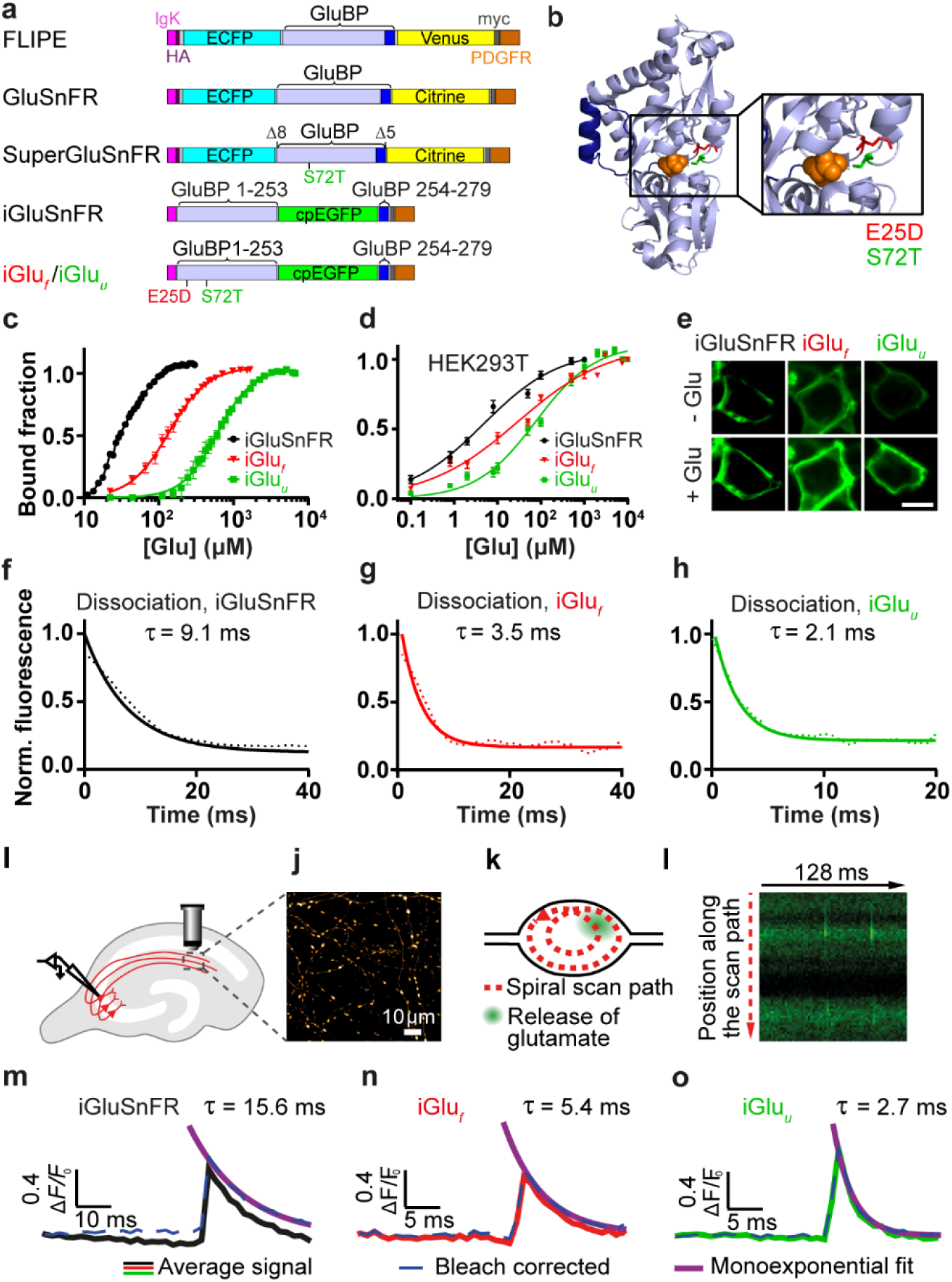
Genetically encoded glutamate indicators (GEGI). (**a**) Domain structure and design of FRET- and single fluorophore-based GEGI; key: (GluBP) (blue), cpEGFP (green), IgG kappa secretion tag (pink), hemagglutinin (HA) tag (purple), myc tag (grey) and a PDGFR transmembrane domain (brown); GluBP 1-253 and 254-279 fragments are in light and dark blue, respectively; Δ8 aa and Δ5 aa specify deletions at the N- and C-terminus of GluBP introduced in GluSnFR. (**b**) Design of selected iGluSnFR variants. Crystal structure of GluBP (PDB 2VHA, adapted from (8). Selected mutated residues around the glutamate site are shown as red and green backbone. Bound glutamate is represented in orange space filling display. (**c**) Equilibrium glutamate binding titrations at 20 °C for iGluSnFR (•), iGluSnFR E25D (iGlu_*f*_) (▾) and iGluSnFR S72T (iGlu_*u*_) (▪) *in vitro*; (**d**) Glutamate titrations *in situ* at 37 °C. iGluSnFR, iGlu_*f*_ and iGlu_*u*_ were expressed in HEK293T cells and titrated with glutamate. Data derived from iGluSnFR (n = 19), iGlu_*f*_ (n = 41) and iGlu_*u*_ (n = 33). (**e**) Representative images of HEK293T cells prior to glutamate addition and at saturating (1, 3 and 10 mM, respectively) glutamate. The scale bar represents 10 μm. Glutamate dissociation kinetics of (**f**) iGluSnFR, (**g**) iGlu_*f*_ and (**h**) iGlu_*u*_ determined by stopped-flow fluorimetry. Experimental data (dotted lines) are overlaid by curves fitted to single exponentials (solid lines). Fluorescence changes are normalised to *F*_max_ of 1. Imaging glutamate release from single presynaptic terminals. (**i**) Schematic representation of organotypic hippocampal slice culture with transfected and patch-clamped CA3 pyramidal cell; (**j**) Imaging axonal projections in CA1 (two-photon stack, maximum intensity projection); (**k**) Individual bouton with spiral scan path for 500 Hz sampling; (**l**) Unfolded scan lines (64 lines, 2 ms/line), single trial. The scan line intersected the fusion site of the vesicle in two positions. Decay time (*τ*_off_) measurements with bleach correction (solid lines) for individual experiments by single exponential fit for (**m**) iGluSnFR (n = 13 boutons, 500 Hz sampling rate) and variants (**n**) iGlu_*f*_ (n = 7 boutons, 1 kHz sampling rate) and (**o**) iGlu_*u*_ (n = 7 boutons, 1 kHz sampling rate).

iGluSnFR has high glutamate affinity and a large dynamic range, but reacts relatively slowly. Its fluorescence response is reported to have a decay half-time (*t*_1/2_) of 92 ms upon synaptic glutamate release (12). Imaging iGluSnFR in cultured hippocampal neurons during 10 Hz stimulation shows summation, which, without deconvolution, might indicate that glutamate accumulates during stimulation (15). Deconvolution of the data suggests that glutamate is cleared between release events (15). iGluSnFR itself is too slow for accurate tracking of synaptic glutamate dynamics during high frequency transmission. Here we introduce two fast iGluSnFR variants, iGlu_*f*_ (for ‘fast’) and iGlu_*u*_ (for ‘ultrafast’) and identify the rate-limiting step leading to bright fluorescence upon glutamate binding. In organotypic slice cultures of rat hippocampus, iGlu_*u*_ directly reports discrete synaptic glutamate release events at 100 Hz. Combining high-speed two-photon imaging and electrophysiology, we show that short-term depression of Schaffer collateral AMPA responses is fully accounted for by the depression of glutamate release. Furthermore, we show a tight correlation between paired-pulse facilitation and rapid recovery from post-tetanic depression at individual boutons, suggesting that differential use of presynaptic resources (readily releasable vesicles) determines the filtering properties of CA3 pyramidal cell boutons.

## RESULTS

### Affinity variants of iGluSnFR by binding site mutations

Six variants of iGluSnFR were generated in which residues coordinating glutamate or in the vicinity of the binding site were mutated (9). The variants had a broad range of glutamate affinities with varied fluorescence dynamic ranges. Two of the mutations lowered, and four increased, the *K*_d_ for glutamate. Variants in order of increasing *K*_d_, covering the range of 19 μM to 12 mM, were E25A<E25R<iGluSnFR<E25D<S72T<R24K<T92A, with Hill coefficients in the range of 1.3 to 2.6. iGluSnFR E25A and E25R had dynamic ranges ∼3 and ∼2 with lower *K*_d_ values compared to iGluSnFR whereas the *K*_d_-s for the R24K and T92A variants were in the mM range but with dynamic ranges of ∼ 2 (**SI Appendix Fig. S1a** and **Table S1**).

We selected the two variants with the fastest response kinetics, iGluSnFR E25D (termed iGlu_*f*_) and iGluSnFR S72T (termed iGlu_*u*_) (**Fig. 1a,b**) for detailed characterization with regard to their biophysical properties as isolated proteins and as membrane-bound glutamate sensors on HEK293T cells and pyramidal neurons. Selectivity for glutamate was determined against aspartate, glutamine, D-serine, GABA and glycine. iGlu_*f*_ and iGlu_*u*_ affinities for aspartate were similar to that for glutamate, as previously reported for iGluSnFR (Marvin et al., 2013). The fluorescence enhancement evoked by aspartate binding was however 2 to 3-fold reduced compared to that by glutamate. The affinity for glutamine was in the mM range for all three probes (**SI Appendix Fig. S1b, Table S3**). D-serine, GABA and glycine evoked no detectable response. p*K*_a_for the glutamate-bound form was ∼6.5 for iGluSnFR iGlu_*f*_ and iGlu_*u*_, whereas the apo-form showed little pH dependence indicating a well-shielded chromophore (**SI Appendix Fig. S1c-e**). Brightness values for iGlu_*f*_ and iGlu_*u*_ were similar to that for iGluSnFR (**SI Appendix Table S2**). *In vitro* measurements gave a dissociation constant (*K*_d_) for glutamate of 33 μM for iGluSnFR, similar to that previously reported (12), while iGlu_*f*_ and iGlu_*u*_ had increased *K*_d_ values of 137 μM and 600 μM, respectively (**Fig. 1c, SI Appendix Table S1**). When expressed on the membrane of HEK293T cells, *K*_d_ values for glutamate were reduced to 3.1 ± 0.3 μM for iGluSnFR, 26 ± 2 μM for iGlu_*f*_ and 53 ± 4 μM for iGlu_*u*_ (measured at 37 °C, **Fig. 1d,e**). A similar reduction of the *K*_d_ in the cellular environment compared to that in solution was reported for iGluSnFR (12). The *in situ* fluorescence dynamic range (*F*_+Glu_ - *F*_-Glu_)/*F*_-Glu_ orΔ*F*/*F*_0_) was 1.0 ± 0.1 for both iGluSnFR and iGlu_*f*_, but 1.7-fold larger for iGlu_*u*_.

### Kinetic measurements of iGluSnFR variants in vitro and in situ

Based on their large *K*_d_ values, we expected iGlu_*f*_ and iGlu_*u*_ to have faster glutamate release kinetics than iGluSnFR. Fluorescence measurements in a stopped-flow instrument indeed revealed faster *off*-rates for the new variants: using the non-fluorescent high-affinity GluBP 600n (10) in excess (0.67 mM) to trap released glutamate, *k*_off_ values of 110 s-^1^ (*τ*_off_ = 9 ms), 283 s-^1^ (*τ*_off_ = 4 ms) and 468 s-^1^ (*τ*_off_ = 2 ms) were obtained for iGluSnFR, iGlu_*f*_ and iGlu_*u*_, respectively, at 20 °C (**Fig. 1f-h** and **SI Appendix Table S4**). To compare *in vitro* response kinetics to physiological measurements, the temperature dependencies of the *off*-rates of iGluSnFR and the fast variants were determined. Linear Arrhenius plots were obtained between 4 °C and 34 °C (**SI Appendix Fig. S1f,g**). For the fast variants, values exceeding the temporal precision of our stopped-flow device were linearly extrapolated (dotted line in **SI Appendix Fig S1f,g**). At 34 °C, decay rates were 233 ± 3 s-^1^ for iGluSnFR (*τ*_off_ = 4.3 ms), 478 ± 5 s-^1^ for iGlu_*f*_ (*τ*_off_ = 2.1 ms) and 1481 ± 74 s-^1^ iGlu_*u*_ (*τ*_off_ = 0.68 ms). Thus, we were able to improve iGluSnFR kinetics by a factor of 6.3.

To image glutamate dynamics in the synaptic cleft, we expressed the newly generated iGluSnFR variants in CA3 pyramidal cells in organotypic slice culture of rat hippocampus (**Fig. 1i, j**). Fluorescence was monitored at single Schaffer collateral terminals in CA1 by spiral line scanning (**Fig. 1k**) while action potentials were triggered by brief (2 ms) depolarizing current injections into the soma of the transfected CA3 neuron. Glutamate release was detected as a sharp increase in green fluorescence (**Fig. 1l**). The iGluSnFR response started 4.5 ± 1.6 ms (mean ± SD) after the peak of the somatic action potential, consistent with a short propagation delay between CA3 and CA1. Consistent with the stochastic nature of glutamate release, individual boutons showed different release probabilities (median *p*_r_ = 0.56, range 0.05 – 1.0). For kinetic analysis, boutons with high release probability and good separation between release failures and successes were selected (**SI Appendix Fig. S2**). The measured fluorescence decay time constants (*τ*_off_) were 13.8 ± 3.8 ms for iGluSnFR, 5.2 ± 2.0 ms for iGlu_*f*_, and 2.6 ± 1.0 ms for iGlu_*u*_ (**Fig. 1m-o, SI Appendix Fig. S3**). Thus, compared to iGluSnFR, synaptic responses detected by iGlu_*u*_ were revealed to be faster by a factor of 5.3. Interestingly, blocking glutamate uptake with TBOA (40 µM) did not slow down the decay of iGlu_*u*_ fluorescence (**SI Appendix Fig. S4**), suggesting that after sparse activation of Schaffer collateral synapses, glutamate is rapidly cleared from the synaptic cleft by diffusion, not by active transport. The situation may be different in highly active neuropil (12, 16).

### Synaptic glutamate dynamics during high frequency stimulation

With decay kinetics of 1-2 milliseconds, iGlu_*f*_ and iGlu_*u*_ are promising tools for direct tracking of synaptic glutamate during high frequency stimulation. The response of iGluSnFR, iGlu_*f*_ and iGlu_*u*_ to paired-pulse stimulation (**Fig. 2** and **SI Appendix Fig. S2**) and to trains of 10 action potentials (APs) at 50, 67 and 100 Hz (**SI Appendix Fig. S5**) was tested. While the responses of iGluSnFR and iGlu_*f*_ suggested build-up of glutamate during high frequency stimulation, iGlu_*u*_ responses revealed that even at 100 Hz stimulation, glutamate was completely cleared from the synaptic cleft between action potentials (**Fig. 2f, SI Appendix Fig. S5i**). Interestingly, the amplitudes of synaptic fluorescence signals (Δ*F*/*F*_0_) were similar for all three indicators, suggesting that the *on*-rate, not the overall affinity, determined the number of glutamate-bound indicator molecules in the synaptic cleft.

**FIGURE 2.**
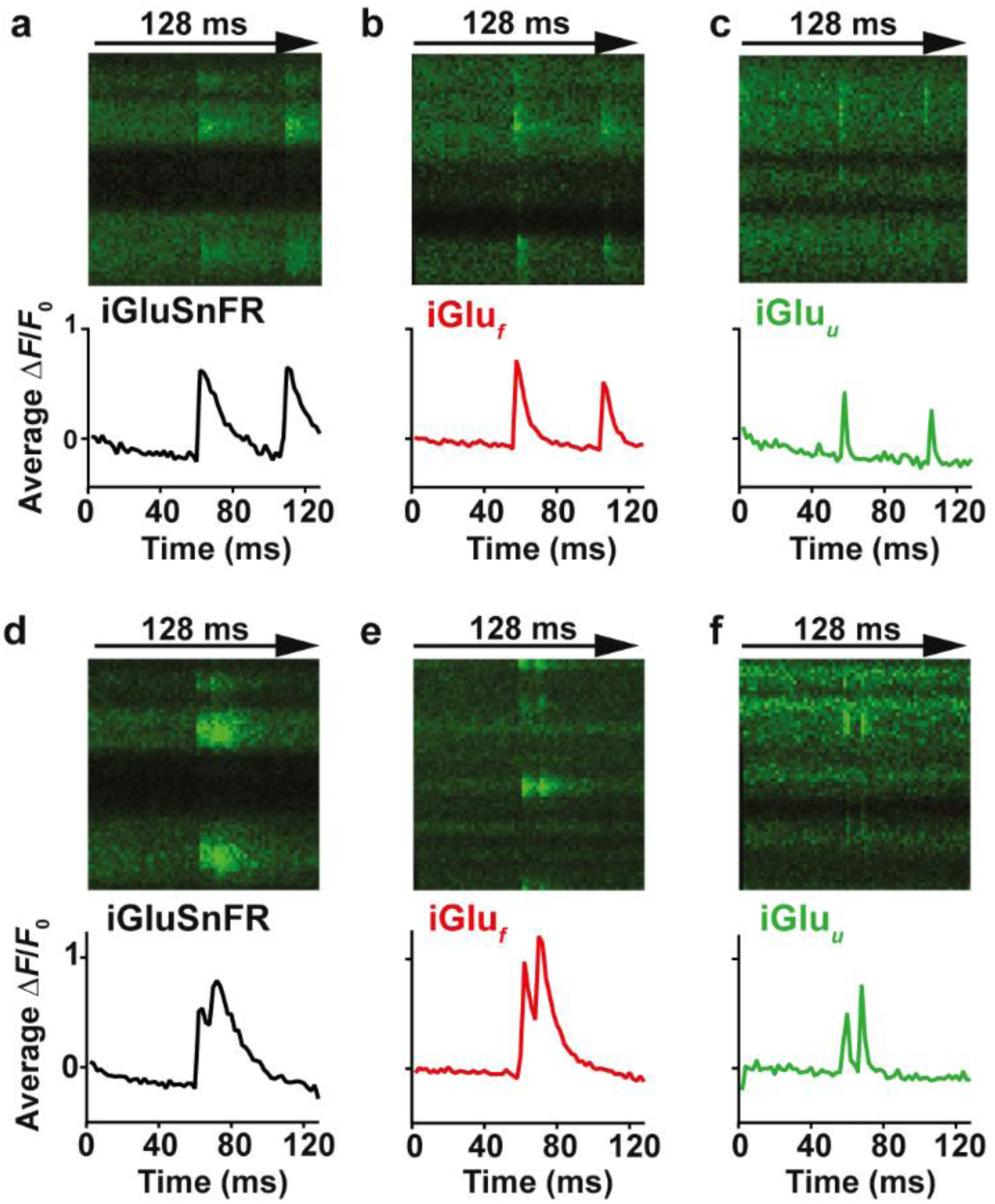
Imaging glutamate release from single presynaptic terminals. Spiral line scans at 500 Hz were used to cover the entire surface of individual boutons, intersecting the release site multiple times. Averages of 3-6 responses of (**a, d**) iGluSnFR, (**b, e**) iGlu_*f*_ and (**c, f**) iGlu_*u*_-expressing boutons stimulated by 2 somatic APs at 48 ms (**a-c**) and 10 ms inter-stimulus intervals (**d-f**).

Excitatory postsynaptic potentials (EPSPs) in CA1 become strongly depressed during high-frequency stimulation (17). We were interested whether EPSP depression during 100 Hz stimulation could be fully accounted for by depression of glutamate release from presynaptic boutons. In paired recordings from connected CA3-CA1 pyramidal cells, we triggered APs in the CA3 cell by brief current injections while monitoring postsynaptic potentials (EPSPs) in the CA1 cell. The protocol consisted of a short high frequency burst (10 APs at 100 Hz) followed by a single AP 500 ms after the burst to probe recovery of synaptic function (18). We repeated the protocol up to 100 times at 0.1 Hz and averaged the recorded traces (**Fig. 3a**). The decay time constant of the recovery response was used to extract the amplitude of individual responses during the 100 Hz train by deconvolution (**Fig. 3b,d**). As expected, connected CA3-CA1 pyramidal cell pairs showed strong depression during the high frequency train. The response to the recovery test pulse (#11) was not significantly different from the first EPSP in the train, indicating full recovery of synaptic function. To investigate depression and recovery of glutamate release, we evaluated iGlu_*u*_ signals during identical stimulation (**Fig. 3c,e**). Due to the extremely fast kinetics of the indicator, deconvolution of the fluorescence time course was not necessary: We read the peak amplitudes during the 100 Hz train directly from the averaged fluorescence time course (average of 10 individual trials sampled at 1 kHz, **SI Appendix Fig. S6**). Glutamate release decreased during the train with a time course that matched EPSP depression (**Fig. 3c**). This result points to a purely presynaptic origin of depression, which is consistent with AMPA receptors rapidly recovering from desensitization after each release event (r_ecovery_ = 5 ms (19)). However, glutamate release 500 ms after the tetanus was still significantly depressed (two-tailed student’s test, *p*-value: 0.0034) while AMPA receptor currents were not. This discrepancy suggests that the response of AMPA receptors to cleft glutamate was in fact potentiated 500 ms after the high frequency train, compensating for the reduced output of Schaffer collateral boutons.

**FIGURE 3.**
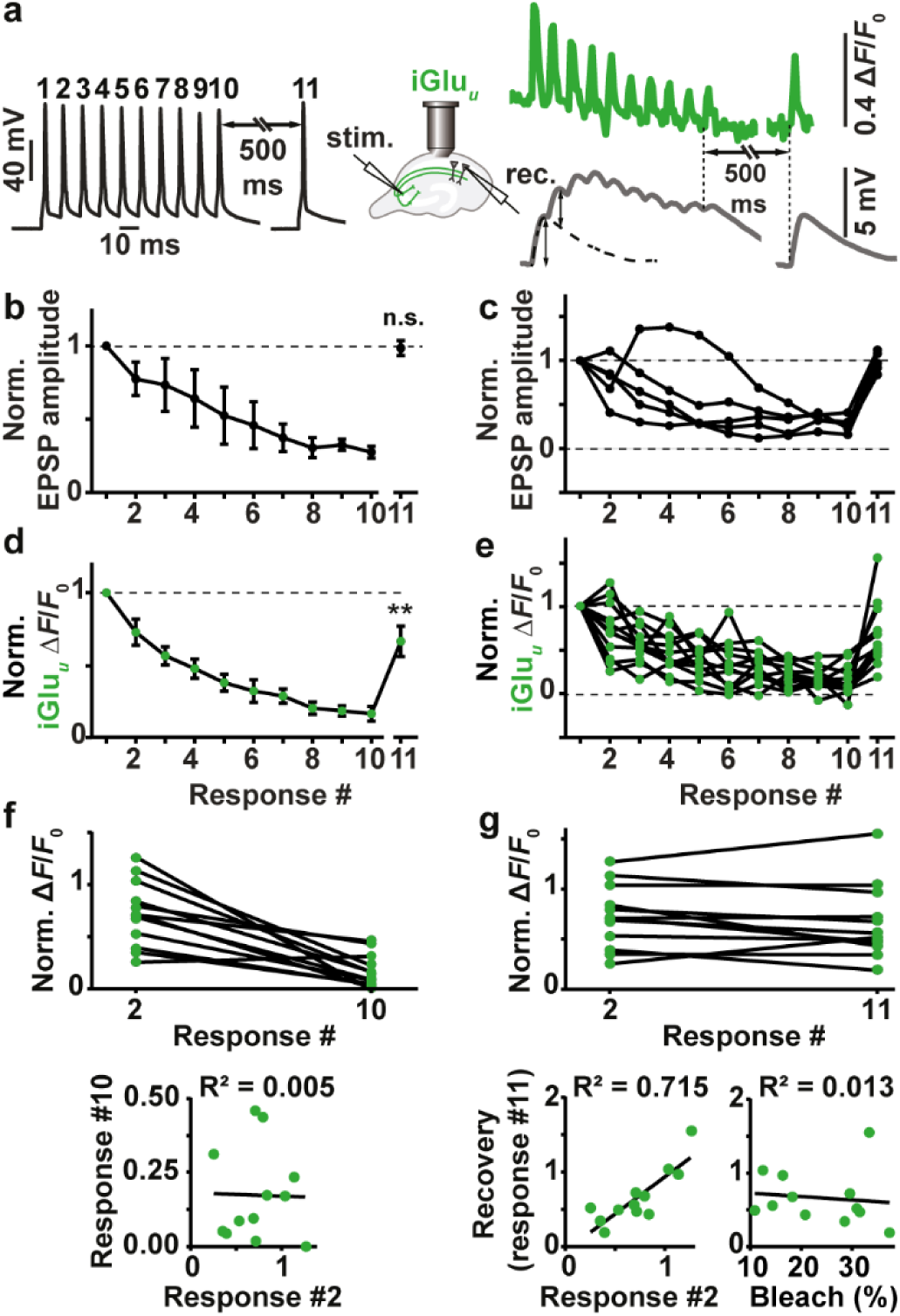
Depression and recovery of synaptic transmission during 100 Hz trains. (**a**) Example of patch-clamp recording from a connected pair of CA3-CA1 pyramidal cells. Black trace: induced action potentials (APs) in CA1 pyramidal cell, 100 Hz train and single AP. Gray trace: EPSPs in CA1 pyramidal cell (average of 50 sweeps). The single AP response (right) was used to extract EPSP amplitudes from the burst response (dotted line). Green trace: average of 10 sweeps of single-bouton iGlu_*u*_ responses to identical stimulation. (**b**) EPSPs (deconvoluted amplitudes) show strong depression during the 100 Hz train, followed by full recovery 500 ms later (n = 5 CA3-CA1 pairs); two-tailed student’s test comparing EPSP #1 and EPSP #11. (**c**) Individual paired recordings show consistent depression (response #10) and recovery (response #11). (**d**) Glutamate release shows strong depression during the 100 Hz train and partial recovery 500 ms later (n = 12 boutons, 8 cells); two-tailed student’s test comparing response #1 and response #11 (p<0.01). (**e**) Individual Schaffer collateral boutons show large variability in response #2 and in recovery response (#11) (**f**) responses by iGlu_*u*_ to the second AP (paired-pulse facilitation/depression) were not correlated with total depression (response #10 normalized to response #1). (**g**) iGlu_*u*_ responses to the second AP (response #2 normalized to response #1) were highly correlated with recovery after 500 ms (response #11 normalized to response #1). Recovery was independent of indicator bleach (*F*_0, response #11_/ *F*_0, response #1_).

### Paired-pulse facilitation correlates with rapid recovery from depression

The rapid kinetics of iGlu_*u*_ allowed us to analyze frequency filtering at individual boutons. On the second AP, boutons showed a wide range of facilitated (3 out of 12 boutons) or depressed responses (9 out of 12 boutons, **Fig. 3e**). The response to the tenth AP was strongly depressed in all boutons (16% of response amplitude to first AP), with no correlation between the second and the tenth response (R_2_ = 0.005, **Fig. 3f**). Interestingly, a highly significant correlation was observed between the response to the second AP and the recovery response 500 ms after the high frequency train (R_2_ = 0.72, **Fig. 3g**). In contrast, we found no correlation between the amount of bleaching in individual experiments (*F*_0_ before 11th pulse / *F*_0_ before 1st pulse) and the amplitude of the recovery response ((Δ*F*/*F*_0_)_11th pulse_ / (Δ*F*/*F*_0_)_1st pulse_), indicating that poor recovery was not caused by excessive bleaching or dilution of indicator molecules. Thus, synapses that showed pronounced paired-pulse facilitation were also able to recover rapidly from depression, both of which is indicative of a low utilization of presynaptic resources (18). Such boutons are optimized for the transmission of high-frequency activity (spike bursts). In contrast, boutons that showed paired-pulse depression were still depressed 500 ms after the high-frequency train. These boutons act as low-pass filters: they preferentially transmit isolated APs preceded by a silent period.

### Response kinetics of iGluSnFR and variants iGlu_*f*_ and iGlu_*u*_ are based on the rate of structural change

Finally, we investigated the response mechanism of iGluSnFR and its fast variants using fluorescence stopped-flow with millisecond time resolution. In association kinetic experiments (20 °C), the fluorescence response rates (*k*_obs_) showed hyperbolic glutamate concentration dependence, approaching saturating rates of 643 s^−1^ and 1240 s^−1^ for iGluSnFR and iGlu_*f*_, respectively (**Fig. 4a-d**). For iGlu_*u*_, in contrast, *k*_obs_ was found to be concentration-independent at 604 s^−1^ (**Fig. 4e,f**). We considered two different reaction pathways to explain our kinetic data (**Fig. 4g)**. iGluSnFR is represented as a complex of the large fragment of the GluBP domain (GluBP 1-253, iGlu_l_), N-terminally flanking cpEGFP and of the C-terminally fused small GluBP fragment (GluBP 254-279, iGlu_s_). The term iGlu_l_∼iGlu_s_, indicates that the large GluBP fragment iGlu_l_ and the small fragment iGlu_s_ are within one molecule, albeit separated by the interjecting cpEGFP. In **Scheme 1**, the binding of glutamate to iGlu_l_ in iGlu_l_∼iGlu_s_ is the primary step (no change in fluorescence). Glutamate binding is followed by a conformational change induced by the reattachment of iGlu_s_ to Glu-bound iGlu_l_, resulting in the highly fluorescent Glu.iGlu_c*_ complex (rate limiting step). According to **Scheme 1**, the hyperbolic dependence of the observed rate *k*_obs_ on the glutamate concentration [Glu] has the intercept of the y-axis at *k*_-2_ (see **SI Appendix Kinetic Theory, eq. 7**). At low [Glu], the initial linear slope gives *k*_+2_*K*_1_. At high [Glu], *k*_obs_ tends to *k*_+2_+*k*_-2_. Although *k*_obs_for iGlu_*u*_ appears essentially concentration independent, its kinetics is consistent with **Scheme 1**, with *k*_+2_+*k*_-2_ having a similar value to *k*_-2_ (**SI Appendix Table S5)**.

**FIGURE 4.**
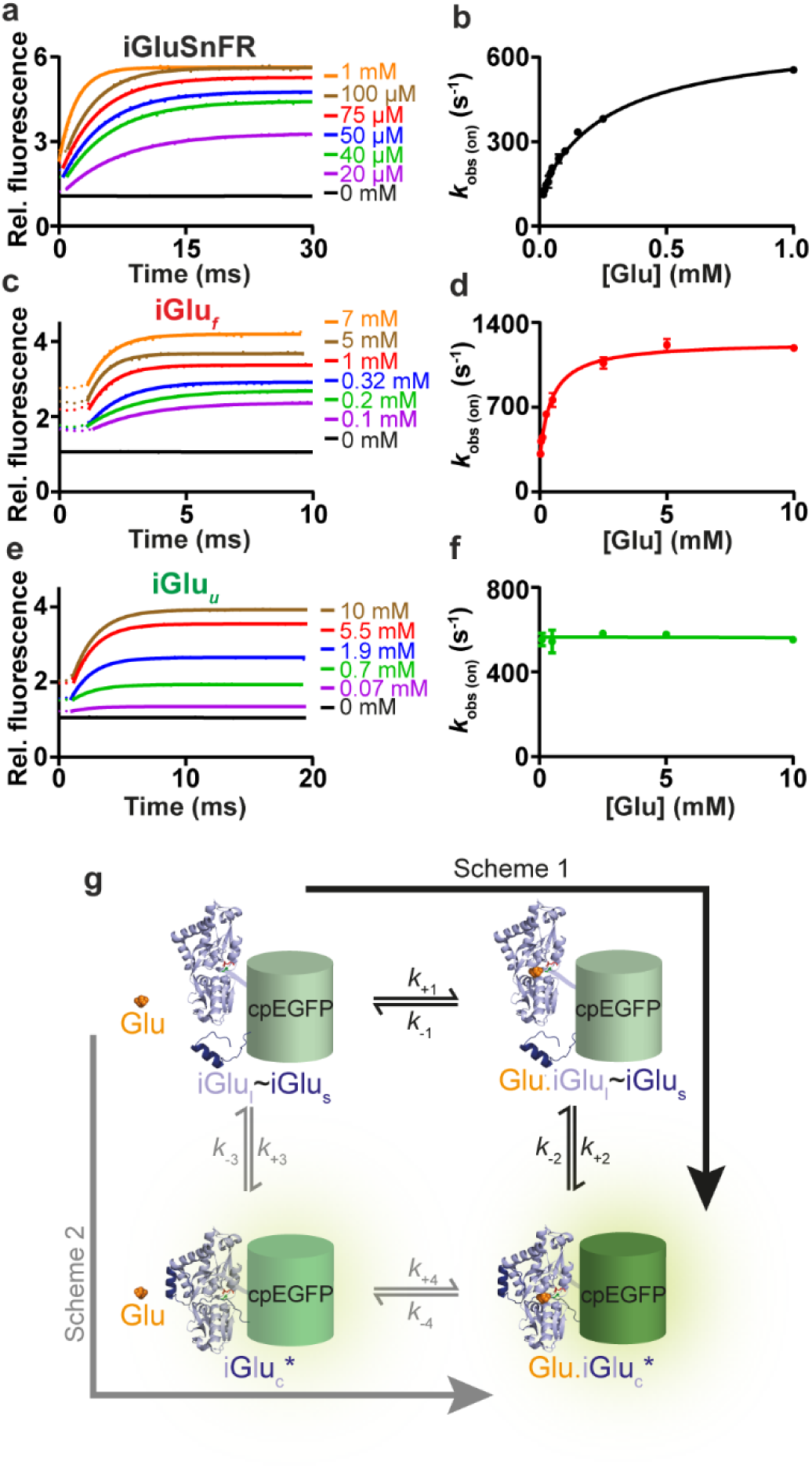
Kinetics of glutamate binding by iGluSnFR variants (20 °C). (**a, c, e**) Glutamate association kinetics of iGluSnFR, iGlu_*f*_ and iGlu_*u*_, respectively. Stopped-flow records of iGluSnFR, iGlu_*f*_ and iGlu_*u*_ reacting with the indicated concentrations of glutamate. Experimental data (dotted lines) are overlaid with curves fitted to single exponentials (solid lines); (**b, d, f**) Plot of observed association rates, *k*_obs(on)_ of iGluSnFR, iGlu_*f*_ and iGlu_*u*_ as a function of glutamate concentration; (**g**) Cartoon diagram depicting the putative molecular transitions of iGluSnFR and its fast variants to the fluorescent state. Key: cpEGFP (green), GluBP 1-253 (iGlu_l_) (light blue) and 254-279 (iGlu_s_) (dark blue) fragments, glutamate (orange).

In the alternative pathway (**Scheme 2)**, the reattachment of iGlu_s_ to iGlu_l_ occurs without prior binding of glutamate. Therefore, iGlu_l_∼iGlu_s_ with the GluBP fragments separated and complete GluBP domain (iGlu_c_*) are in equilibrium. The conformational change that represents the reattachment of the two GluBP fragments is expected to generate a fluorescent state of cpEGFP. However, the equilibrium is likely to be strongly shifted to the separated, non-fluorescent state (iGlu_l_∼iGlu_s_). Assuming that this equilibrium is fast and glutamate binding stabilizes the fluorescent state, at low [Glu], a linear dependence of *k*_obs_ on [Glu] is predicted with a slope of *K*_3_*k*_+4_/(1+*K*_3_) and an intercept of the y-axis at *k*_-2_ (see **SI Appendix Kinetic Theory, eq. 15)**. Although at low [Glu], mono-exponential fluorescence changes are expected, as [Glu] increases, the concentration of iGlu_c_* cannot be assumed to be at steady-state and slow isomerisation will limit *k*_obs_, in a similar pattern to that for Scheme 1. Thus, at high [Glu], even if iGlu_c_* and Glu.iGlu_c_* have equal relative fluorescence intensities, biphasic fluorescence changes would be expected for the association reactions. As all the reactions studied here for the three variants had a single exponential appearance, we can exclude **Scheme 2** as a possible reaction pathway. In conclusion, **Scheme 1** provides an excellent fit to our measurements (**SI Appendix Table S5)**, pointing to ‘Venus fly-trap’ closure by glutamate binding as a required first step for the conformational change that increases iGluSnFR fluorescence.

## DISCUSSION

The development of iGluSnFR was a breakthrough in fluorescent glutamate sensors towards investigating neurotransmission in living organisms (20). Here we describe how to overcome one of the key limitations of iGluSnFR, its slow response kinetics, and use the new utrafast variant iGlu_*u*_ to investigate synaptic transmission and frequency filtering at individual Schaffer collateral boutons.

For all tested variants, synaptic *off*-kinetics were slower by a factor of 2.5 - 3.8 compared to temperature-matched *in vitro* measurements on isolated protein. This is consistent with the much higher affinities of HEK293T cell-expressed glutamate sensors compared to soluble protein. These systematic differences, also noted in the original characterization of iGluSnFR (12), may be attributed to the tethering of the molecule to a membrane anchor, slowing down conformational changes compared to free-floating sensor molecules. Nevertheless, the relative differences in affinity and kinetics of the new versions compared to iGluSnFR were preserved *in vitro* and *in situ*. The *on*- and *off*-rates of iGlu_*u*_ are greater (2- and 5-6 fold, respectively) compared to iGluSnFR. Interestingly, iGlu_*u*_ was a faster reporter in the hippocampal slice than iGlu_*f*_, even though the latter has a faster limiting *on*-rate. iGlu_*u*_ may be put at an advantage over iGlu_*f*_ by its concentration-independent response kinetics. It must be noted that the kinetics of iGluSnFR-type indicators are ultimately limited by the structural change that reconstitutes the fluorescent complex, similar to calcium-sensing GCaMPs. The constraints of the mechanism with regard to the onset of fluorescence suggest that it cannot be engineered to resolve sub-millisecond glutamate dynamics. To achieve microsecond response times, it might be necessary to develop hybrid glutamate indicators using synthetic dyes.

Synaptic iGlu_*u*_ imaging revealed complete clearance of glutamate between release events even at 100 Hz stimulation frequency. The first attempts to estimate the time course of synaptic glutamate transients were based on the decay of NMDA receptor responses in primary cell culture: kinetic analysis of the displacement of a competitive NMDA receptor antagonist suggested glutamate clearance with τ = 1.2 ms (21). More recent studies using computational modeling and fluorescence anisotropy imaging in tissue suggest that it is closer to 100 μs (22, 23). Thus, due to the intrinsic kinetic limits of the iGluSnFR mechanism, even iGlu_*u*_ cannot resolve the true dynamics of free glutamate in the synaptic cleft. What we can say with confidence is that accumulation of glutamate in the synaptic cleft does not contribute to short-term plasticity at Schaffer collateral synapses.

In the analysis of synaptic responses, we did not correct for the non-linearity of the iGluSnFR variants (**Fig. 1c**), as response amplitudes (Δ*F*/*F*_0_ of 0.4 – 1.2) were below the *K*_d_ for glutamate (**SI Appendix Table S3**). Therefore, we could use the relative fluorescence changes as a rough estimate of cleft glutamate. Glutamate release showed strong depression during 100 Hz firing, in line with the expected depletion of release-ready vesicles. As we controlled the generation of every action potential by somatic current injections, we can exclude decreased afferent excitability as a source of depression in these experiments (17). AMPA receptor currents during 100 Hz firing did not show more run-down than iGlu_*u*_ responses, suggesting that AMPA receptor desensitization did not play a major role in the decrease of synaptic efficacy during the train. Paradoxically, AMPA responses were fully recovered 500 ms after the train while the iGlu_*u*_ response was still significantly depressed. The most parsimonious explanation is a long-lasting depression of glutamate release. There are alternative scenarios that could explain smaller iGlu_*u*_ responses on the 11th pulse, e.g. indicator molecules retrieved into endosomal structures during endocytosis, or accumulation of indicator in a (hypothetical) desensitized state. In these scenarios, facilitating boutons, which experience more exo- and endocytosis and iGlu_*u*_ activation during the train, would be expected to show smaller responses at the 11_th_ pulse. However, we found a strong correlation in the opposite direction, making these scenarios less likely (**Fig. 3g**).

The full recovery of the AMPA response points to an unexpected increase in sensitivity of the postsynaptic compartment to glutamate. By association with different auxiliary proteins and other scaffold-related mechanisms, the density and open probability of postsynaptic glutamate receptors can quickly change (24, 25). In hippocampal slice cultures, post-tetanic potentiation is well established and requires the activity of protein kinase C (26). Thus, it is possible that elevated Ca_2+_ levels in the spine during our high frequency protocol enhanced AMPA receptor currents by a number of mechanisms, compensating for the reduced glutamate release 500 ms after the tetanus.

The surprisingly tight correlation between paired-pulse facilitation and rapid recovery from depression at individual boutons provides direct evidence that differential use of presynaptic resources determines the neural code between pyramidal cells (1, 18). Using Schaffer collateral synapses as an example, we show that iGlu_*u*_ is a useful tool for a mechanistic analysis of high frequency synaptic transmission, interrogating presynaptic function independently of postsynaptic transmitter receptors.

## METHODS

We provide a detailed description of the methods, data analysis and kinetic modeling in the online Supporting Information. Plasmids for iGlu_*f*_, iGlu_*u*_ will be deposited at Addgene and made available once the manuscript is accepted for publication.

### Materials

pCMV(MinDis).iGluSnFR and pRSET FLIPE-600n plasmids were a gift from Loren Looger (Addgene Plasmid #41732) and Wolf Frommer (Addgene plasmid # 13537), respectively. Site-directed mutagenesis was carried out following the QuikChange II XL protocol (Agilent Technologies).

### Fluorescence spectroscopies

Glutamate association and dissociation kinetic experiments of iGluSnFR proteins were carried out on a Hi-Tech Scientific SF-61DX2 stopped-flow system equipped with a temperature manifold (27). Fluorescence spectra and equilibrium glutamate titrations were recorded on a Fluorolog3 (Horiba Scientific).

### *In situ* glutamate titration

HEK293T cells were cultured on 24-well glass bottom plates in DMEM containing non-essential amino-acids (Life Technologies), 10% heat inactivated FBS (Life Technologies) and penicillin/streptomycin (100 U/mL, 100 mg/mL, respectively), at 37 °C in an atmosphere of 5% CO_2_. Cells were allowed 24 h to adhere before transfection with Lipofectamine 2000 (Invitrogen). Cells were examined at 37 °C (OKO lab incubation chamber) with a 3i Marianas spinning-disk confocal microscope equipped with a Zeiss AxioObserver Z1, a 40x/NA1.3 oil immersion objective and a 3i Laserstack as excitation light source (488 nm).

### Synaptic measurements

Organotypic hippocampal slices (400 μm) were prepared from male Wistar rats at postnatal day 5 as described (28). iGluSnFR and variant plasmids were electroporated into 2-3 CA3 pyramidal cells at 40 ng/µL (iGluSnFR) or 50 ng/µL (iGlu_*f*_, iGlu_*u*_) together with tdimer2 (20 ng/µL), a cytoplasmic red fluorescent protein (29). 2 - 4 days after electroporation (at DIV 14-30), slice cultures were placed in the recording chamber of a two-photon microscope and superfused with artificial cerebrospinal fluid (ACSF) containing (in mM) 25 NaHCO_3_, 1.25 NaH_2_PO_4_, 127 NaCl, 25 D-glucose, 2.5 KCl, 2 CaCl_2_, 1 MgCl_2._ Whole-cell recordings from a transfected CA3 pyramidal cell were made with a Multiclamp 700B amplifier (Molecular Devices). Red and green fluorescence was detected through the objective (LUMPLFLN 60XW, 60x, NA 1.0, Olympus) and through the oil immersion condenser (NA 1.4, Olympus) using 2 pairs of photomultiplier tubes (H7422P-40SEL, Hamamatsu).

## Acknowledgements

We thank Dr Zoltan Ujfalusi (University of Kent) for assistance with stopped-flow experiments, Iris Ohmert for the preparation of organotypic cultures and Dr David Trentham for comments on the manuscript. Use of The Institute of Translational Medicine, University of Liverpool Imaging Facility is gratefully acknowledged. This project was supported by the Wellcome Trust (Grant 094385/Z/10/Z to K.T.) and BBSRC (Grant BB/M02556X/1 to K.T.); the German Research Foundation (SPP 1665, SFB 936, FOR 2419 to T.G.O; SPP 1926, FOR 2419 to J.S.W) and the European Research Council (ERC-2016-StG 714762 to J.S.W.).

